# Deletion of Murine Gammaherpesvirus Gene *M2* in AID-Expressing B Cells Impairs Host Colonization and Viral Reactivation

**DOI:** 10.1101/2020.03.30.016899

**Authors:** Shana M. Owens, Darby G. Oldenburg, Douglas W. White, J. Craig Forrest

## Abstract

Gammaherpesviruses (GHVs) are DNA tumor viruses that establish life-long, chronic infections in lymphocytes of humans and other mammals. GHV infections are associated with numerous cancers, especially in immune compromised hosts. While it is known that GHVs utilize host germinal center (GC) B cell responses during latency establishment, an understanding of how viral gene products function in specific B cell subsets to regulate this process is incomplete. Using murine gammaherpesvirus 68 (MHV68) as a small-animal model to define mechanisms of GHV pathogenesis *in vivo*, we generated a virus in which the *M2* gene was flanked by *loxP* sites (M2.loxP), enabling the use of Cre-lox technology to define M2 function in specific cell types in infection and disease. The *M2* gene encodes a protein that is highly expressed in GC B cells that promotes plasma cell differentiation and viral reactivation. *M2* was efficiently deleted in Cre-expressing cells, and the presence of *loxP* sites flanking *M2* did not alter viral replication or latency in mice that do not express Cre. In contrast, M2.loxP MHV68 exhibited a deficit in latency establishment and reactivation that resembled M2-null virus, following intranasal (IN) infection of mice that express Cre in all B cells (CD19-Cre). Nearly identical phenotypes were observed for M2.loxP MHV68 in mice that express Cre in germinal center (GC) B cells (AID-Cre). However, neither colonization of draining lymph nodes after IN infection nor the spleen after intraperitoneal (IP) infection required M2, although the reactivation defect was retained. Together, these data confirm that M2 function is B cell-specific and demonstrate that M2 primarily functions in AID-expressing cells to facilitate MHV68 dissemination to distal latency reservoirs within the host and reactivation from latency. Our study reveals that a viral latency gene functions within a distinct subset of cells to facilitate host colonization.

**IMPORTANCE:** Gammaherpesviruses establish life-long chronic infections in cells of the immune system that can lead to lymphomas and other diseases. To facilitate colonization of a host, gammaherpesviruses encode gene products that manipulate processes involved in cellular proliferation and differentiation. Whether and how these viral gene products function in specific cells of the immune system is poorly defined. We report here the use of a viral genetic system that allows for deletion of specific viral genes in discrete populations of cells. We employ this system in an *in vivo* model to demonstrate cell-type-specific requirements for a particular viral gene. Our findings reveal that a viral gene product can function in distinct cellular subsets to direct gammaherpesvirus pathogenesis.

## INTRODUCTION

Gammaherpesviruses (GHVs) are DNA tumor viruses that include the human pathogens Epstein-Barr virus (EBV) and Kaposi sarcoma-associated herpesvirus (KSHV). GHVs are lymphotropic viruses with a biphasic infection cycle that is characterized by two distinct stages: productive lytic replication or chronic, life-long latency (1). Lytic replication is characterized by temporally regulated viral gene expression, replication of viral DNA, and production of infectious viral progeny (1). After resolution of acute infection, latency is established and viral genomes are maintained within germinal center (GC) and memory B cells (2). During latency, viral gene expression is restricted to a few latency gene products and non-coding RNAs (3, 4). These molecules function to manipulate host-cell physiology, maintain viral genomes, and thwart immune surveillance (5-10). In response to specific stimuli, latent viral genomes can reinitiate the lytic cycle, a process called reactivation (11). Reactivation enables infection of new hosts and is thought to help maintain latent reservoirs within a chronically infected host (11, 12).

To define the processes used by GHVs to infect a living organism and cause disease, we and others utilize the natural rodent pathogen murine gammaherpesvirus 68 (MHV68, also known as murid herpesvirus 4 or MuHV-4) (12). The MHV68 genome is co-linear with EBV and KSHV, and like other GHVs, MHV68 preferentially infects B lymphocytes and can cause cancer, especially in immune compromised animals (13, 14). Because MHV68 readily infects and establishes chronic infection in inbred, outbred, and genetically modified mouse strains (11, 15), MHV68 serves as a tractable small animal model for studying the function of viral gene products and the host immune response to GHV infections.

During latency establishment GHVs are thought to augment and usurp the GC reaction (11, 16, 17). The GC reaction is a B cell maturation process that is characterized by rapid cellular proliferation and the accumulation of mutations in the immunoglobulin locus, which encodes the B cell receptor and antibodies, and is mediated by the enzyme activation-induced cytidine deaminase (AID) (18-20). Subversion of the GC response by GHVs is thought to expand the reservoir of latently infected cells and promote the infection of long-lived memory B cells (MBCs) (21). While it is well known that GHVs establish and maintain chronic infection within GC B cells, requirements for specific viral gene products in this process are still being investigated.

The MHV68 *M2* gene is highly expressed in GC B cells and, while M2 has no known viral or cellular homologs at the sequence level, it is hypothesized to be a functional homolog of the EBV latent membrane protein 2A (LMP2A) and the KSHV K1 protein (22). Transcription of *M2* within B cells is driven by multiple closely-linked promoters, which generate two spliced and one unspliced *M2* transcript. The spliced transcripts encode the M2 protein and are apparently regulated in a B cell-specific manner, as spliced transcripts are not detected during lytic infection, whether in cell culture or the lung epithelium (4, 23). The M2 protein functions as a scaffold that interacts with membrane-associated signaling molecules to mimic B cell-receptor activation and promote calcium-mediated activation of the nuclear factor of activated T cells (NFAT) pathway (24). NFAT activation driven by M2 induces the plasma cell-associated transcription factor, interferon regulatory factor 4 (IRF4), which enforces a gene expression program involved in plasma cell differentiation and promotes production of anti-inflammatory cytokines, such as IL-10 (10, 24). Studies using M2-null MHV68 (M2.Stop) and viruses with specific mutations in putative signaling residues demonstrated that M2 facilitates latency establishment in the spleen after IN, but not IP, inoculation of mice (25-27). M2 is also generally required for viral reactivation from latency *in vivo*, a phenotype that correlates with reduced plasma cell infection by M2-null virus (28). Forced retroviral expression of M2 in primary murine B cells in tissue culture and following *in vivo* transfer drives B cells toward a GC B cell phenotype, which ultimately differentiate, into plasma cells (28, 29). While detection of spliced *M2* transcripts in B cells, but not other cell types, suggests a B cell-specific function, distinct requirements for M2 in unique B cell subsets during viral colonization of the host are not defined.

Since GC B cells are critical early targets for GHV infection, determining how specific viral gene products function within these cells to permit latency is fundamental to understanding GHV pathogenesis. We previously reported development of a viral genetic platform that allows for dissection of cell-type-specific roles of viral gene products *in vivo* (30). Building from this technology, we engineered a recombinant MHV68 in which the gene encoding M2 was flanked by *loxP* sequences (floxed; M2.loxP) to enable the conditional deletion of *M2* in cells that express Cre recombinase. We used this system to define the function of M2 in specific cell-types in mice, especially GC B cells, during latency establishment by MHV68.

## RESULTS

### Generation and validation of M2.loxP MHV68

Teasing apart the function of GHV gene products *in vivo* is complicated in part by the biphasic infection cycle, as well as the convoluted dissemination process required to establish splenic latency. Traditional methods of viral mutagenesis result in complete ablation of a gene of interest. This approach discounts the role of multi-functional proteins that are required in different phases of infection or within distinct cell-types. *M2* encodes a latency-specific gene product that facilitates MHV68 establishment of latency and reactivation and promotes B cell differentiation in infected mice (3, 27-29). To better define roles for the MHV68 latency gene product M2 within distinct populations of B lymphocytes, we generated a recombinant virus to enable cell-type-specific deletion of the *M2* gene in infected cells that express Cre recombinase. We previously demonstrated the feasibility and utility of this approach in a study that defined B cell-specific requirements for *ORF73*, the gene that encodes the MHV68 latency-associated nuclear antigen 105 (mLANA) (30).

In the new MHV68 recombinant, *loxP* sites were inserted flanking the *M2* gene within the MHV68 BAC, and viral stocks were generated (M2.loxP MHV68, **Fig 1A**). Appropriate BAC modification was confirmed by restriction fragment length polymorphism analysis (RFLP, **Fig 1B**) and sequencing of the targeted genetic locus. To confirm that floxed *M2* was efficiently deleted in the presence of Cre, we infected either Vero or Vero-Cre cells with WT MHV68, M2.Stop, or M2.loxP, and then evaluated *M2* locus integrity by PCR. Although intact *M2* was readily detected in Vero cells infected with M2.loxP, only the *M2* deletion amplicon was detected in Vero-Cre cells infected with M2.loxP (**Fig 1C**). The full-length *M2* locus was detected in both cell-types infected by WT MHV68 and M2.Stop. Furthermore, an adjacent viral gene, *M3*, remained intact when *M2* was deleted, suggesting that viral gene deletion is correctly targeted and specific (**Fig 1C**). These results demonstrate that Cre-mediated recombination efficiently deletes the *M2* gene from the M2.loxP MHV68 genome in cell culture.

**Figure 1.**
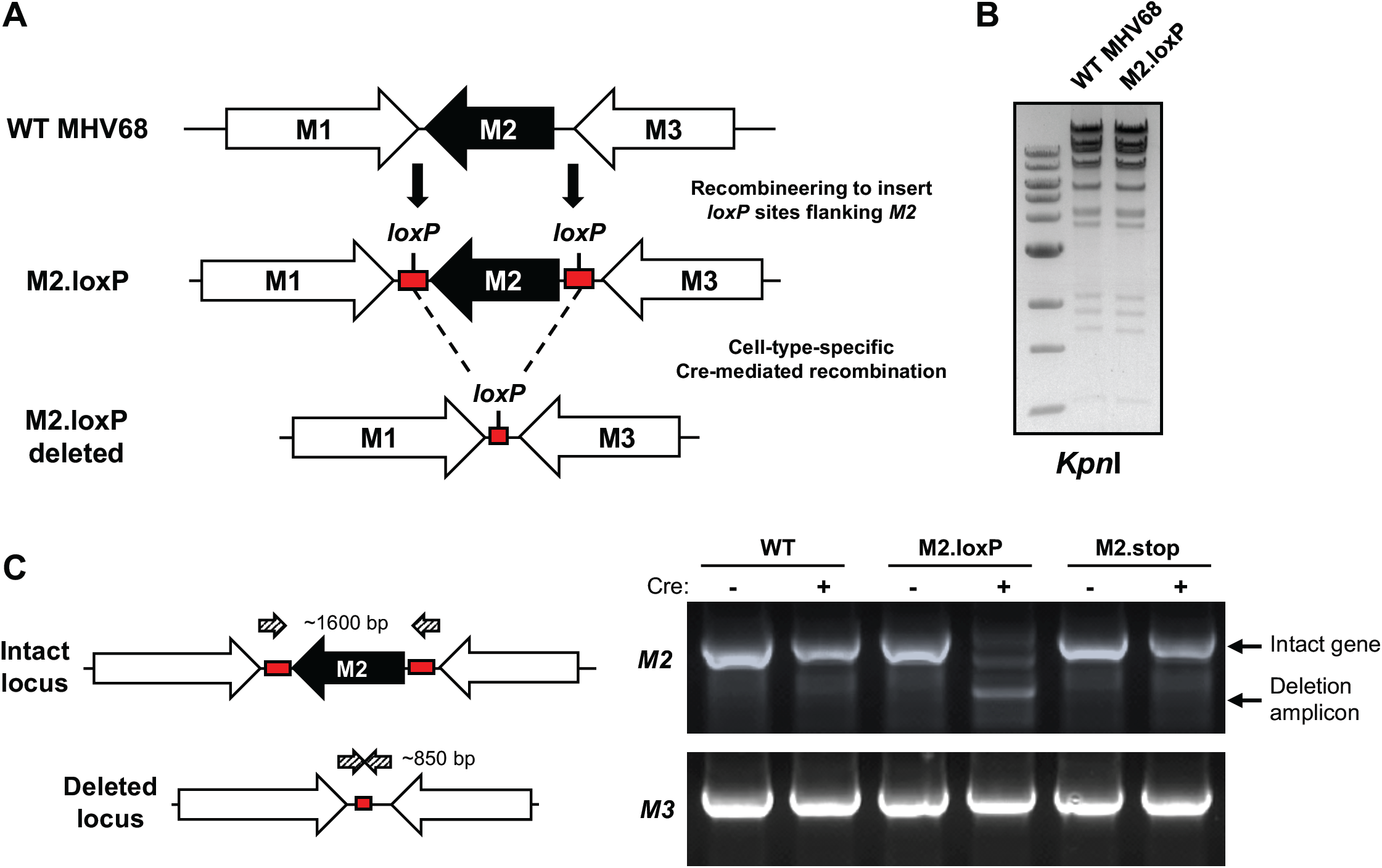
Development and validation of M2.loxP MHV68. (A) Schematic depiction of the insertion of *loxP* sites flanking *M2* in the MHV68 genome and its deletion in the presence of Cre recombinase. (B) BAC DNA was digested with the indicated restriction endonuclease, and digestion products were resolved by agarose-gel electrophoresis to evaluate the gross genetic integrity of the newly derived BAC. (C) Vero or Vero-Cre cells were infected with the indicated viruses at an MOI of 0.05 PFU/cell. Total DNA was isolated on day 4 post-infection, and PCR was performed as illustrated in the schematic to detect the intact or deleted *M2* locus or the adjacent *M3* locus as a control.

To ensure that the presence of *loxP* sites flanking *M2* did not inadvertently attenuate the virus, we evaluated M2.loxP infection relative to control viruses. In cultured cells, M2.loxP MHV68 replicated with similar kinetics to WT MHV68 in both single- and multi-step growth analyses (**Fig 2A**). Further, M2.loxP MHV68 titers were equivalent to WT and M2-null MHV68 on day 7 after IN inoculation in lungs of C57BL/6 mice, a time point that approximates the peak of acute viral replication *in vivo* (**Fig 2B**). This finding confirms that the presence of *loxP* sites did not impact acute viral replication *in vivo*, and M2.Stop results further demonstrate that M2 is not required for lytic replication. However, unlike M2-null virus, M2.loxP established latency in the spleen after IN inoculation of C57BL/6 mice at levels similar to WT MHV68 (**Fig 2C** and **Table 1**). M2.loxP also reactivated from explanted splenocytes efficiently, while M2.Stop was again attenuated (**Fig 2D** and **Table 2**). Together, these data confirm previous findings for M2.Stop and demonstrate that the insertion of *loxP* sites flanking the *M2* gene does not impair MHV68 latency establishment in or reactivation from the spleen of WT mice.

**Table 1.** Frequency of MHV68 latently infected cells.

| Mouse Strain | Virus | Route of infection | Cell population | Day post-infection | Latency frequency |
| --- | --- | --- | --- | --- | --- |
| C57BL/6 | WT MHV68 | IN | Spleen | 16 | 1/500 |
|  | M2.Stop | IN | Spleen | 16 | 1/1500 |
|  | M2.loxP | IN | Spleen | 16 | 1/350 |
| CD19-Cre | WT MHV68 | IN | Spleen | 16 | 1/80 |
|  | M2.loxP | IN | Spleen | 16 | 1/900 |
|  | WT MHV68 | IP | Spleen | 16 | 1/200 |
|  | M2.loxP | IP | Spleen | 16 | 1/300 |
| AID-Cre | WT MHV68 | IN | Spleen | 16 | 1/150 |
|  | M2.loxP | IN | Spleen | 16 | 1/1200 |
|  | WT MHV68 | IP | Spleen | 16 | 1/250 |
|  | M2.loxP | IP | Spleen | 16 | 1/1000 |
|  | WT MHV68 | IP | PEC | 16 | 1/500 |
|  | M2.loxP | IP | PEC | 16 | 1/600 |

**Table 2.**
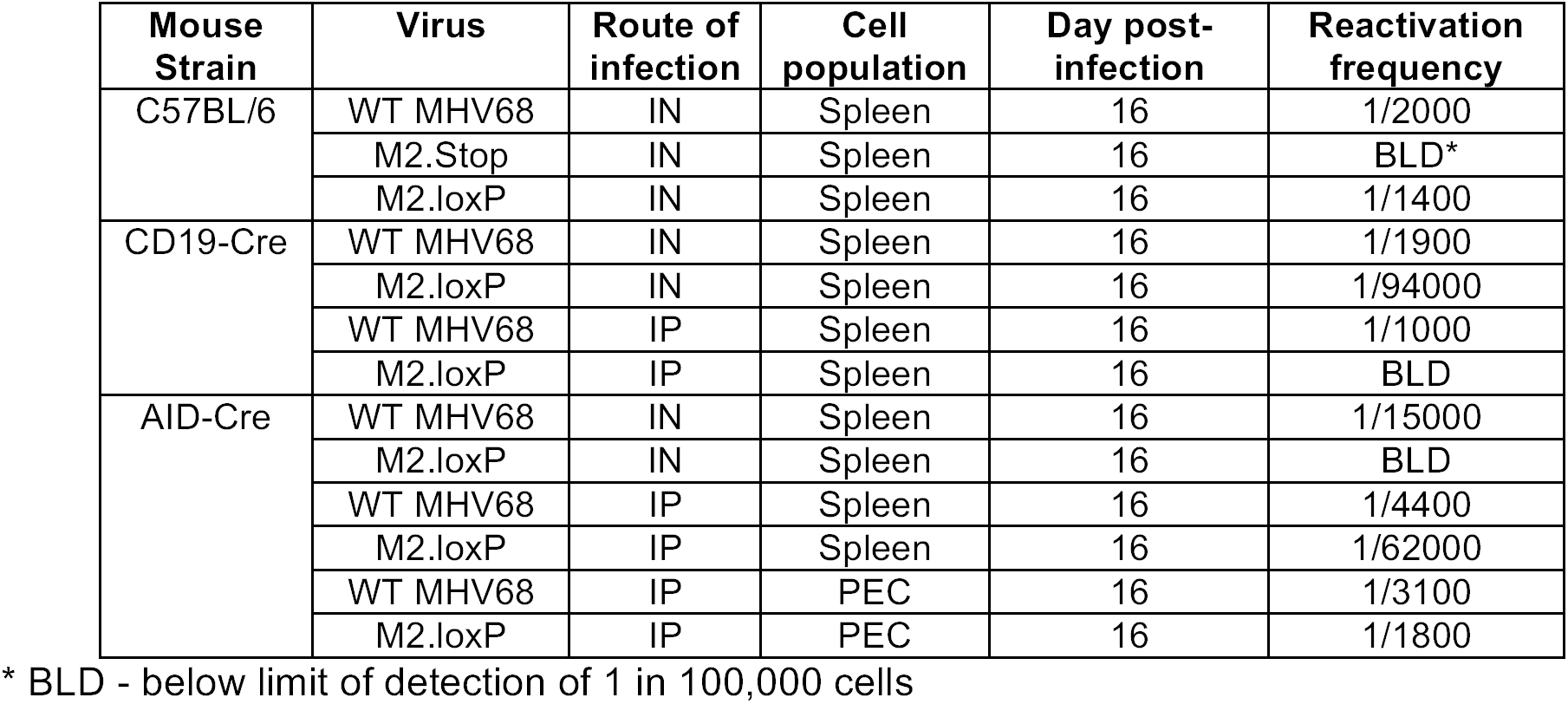
Frequency of reactivation competent cells.

| Mouse Strain | Virus | Route of infection | Cell population | Day post-infection | Reactivation frequency |
| --- | --- | --- | --- | --- | --- |
| C57BL/6 | WT MHV68 | IN | Spleen | 16 | 1/2000 |
|  | M2.Stop | IN | Spleen | 16 | BLD* |
|  | M2.loxP | IN | Spleen | 16 | 1/1400 |
| CD19-Cre | WT MHV68 | IN | Spleen | 16 | 1/1900 |
|  | M2.loxP | IN | Spleen | 16 | 1/94000 |
|  | WT MHV68 | IP | Spleen | 16 | 1/1000 |
|  | M2.loxP | IP | Spleen | 16 | BLD |
| AID-Cre | WT MHV68 | IN | Spleen | 16 | 1/15000 |
|  | M2.loxP | IN | Spleen | 16 | BLD |
|  | WT MHV68 | IP | Spleen | 16 | 1/4400 |
|  | M2.loxP | IP | Spleen | 16 | 1/62000 |
|  | WT MHV68 | IP | PEC | 16 | 1/3100 |
|  | M2.loxP | IP | PEC | 16 | 1/1800 |

\* BLD - below limit of detection of 1 in 100,000 cells

**Figure 2.**
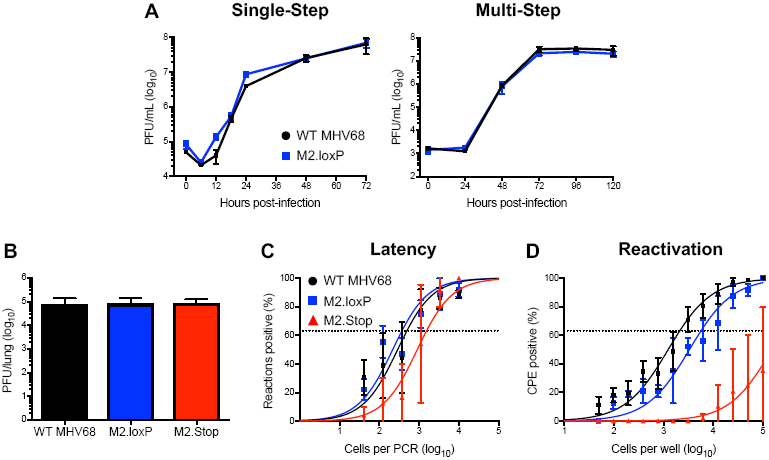
M2.loxP exhibits normal lytic replication and latency in C57BL/6 mice. (A) 3T12 fibroblasts were infected with WT MHV68 or M2.loxP at an MOI of 5 PFU/cell (single-step, left panel) or 0.05 PFU/cell (multi-step, right panel). Viral titers were determined by plaque assay at the indicated times post-infection. Results are means of triplicate samples. Error bars represent standard deviations. (B-D) C57BL/6 mice were infected IN with 1000 PFU of the indicated viruses. (B) Mice were sacrificed on day 7 post-infection, and viral titers in lung homogenates were determined by plaque assay. (C-D) Mice were sacrificed on day 16 post-infection. (C) Single-cell suspensions of spleen cells were serially diluted and the frequencies of cells harboring MHV68 genomes were determined using a limiting-dilution PCR analysis. (D) Reactivation frequencies were determined by *ex vivo* plating of serially diluted cells on an indicator monolayer. Cytopathic effect was scored 2-3 weeks post-plating. Groups of 3–5 mice were pooled for each infection and analysis. Results are means of 2–3 independent infections. Error bars represent standard error of the means.

### M2 is required in CD19^+^ B cells for latency establishment and reactivation

RT-PCR analyses suggest that a properly spliced *M2* transcript capable of yielding the functional M2 protein is only present in B cells (4, 23, 31). From this observation it is hypothesized that M2 primarily functions in B cells to promote MHV68 latency and reactivation. To more conclusively test this hypothesis, we infected mice that express Cre recombinase under control of the B cell-specific CD19 promoter (CD19-Cre) to facilitate *M2* deletion in all B cells. On day 7 after IN inoculation, titers of M2.loxP and WT MHV68 were equivalent in lungs (**Fig 3A**). However, splenomegaly, a consequence of the infectious mononucleosis-like syndrome caused by MHV68, did not occur following infection with M2.loxP (**Fig 3B**). A PCR analysis of the *M2* locus demonstrated that *M2* was deleted in these mice (**Fig 3C**). Latency establishment by M2.loxP also was 10-fold lower than WT MHV68 in spleens of CD19-Cre mice on day 16 post-infection (**Fig 3D** and **Table 1**), and M2.loxP reactivation from explanted splenocytes was ca. 100-fold lower than WT MHV68 (**Fig 3E** and **Table 2**), indicating a further 10-fold reduction over the latency defect. The deficits exhibited by M2.loxP following infection of CD19-Cre mice appear to phenocopy M2.Stop infection of WT C57BL/6 mice (**Fig 2**), indicating that M2 primarily functions in B lymphocytes to facilitate latency establishment and reactivation following IN infection.

**Figure 3.**
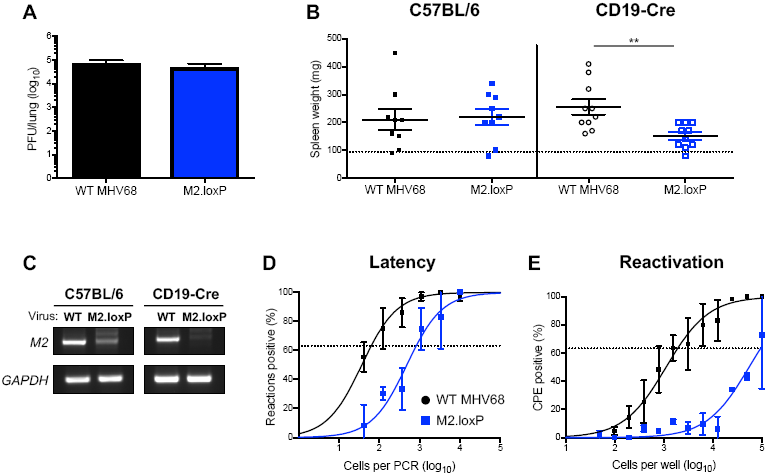
M2 deletion in CD19^+^ cells impairs MHV68 latency establishment and reactivation. CD19-Cre mice were infected IN with 1000 PFU of the indicated viruses. (A) Mice were sacrificed on day 7 post-infection, and viral titers in lung homogenates were determined by plaque assay. (B-E) Mice were sacrificed on day 16 post-infection. (B) Spleens were harvested and weighed as a measure of splenomegaly. The dashed line indicates the average mass of spleens from mock-infected mice. Each dot represents one mouse. Spleen weights from C57BL/6 mice infected in Figure 2 are shown for comparison. (C) DNA was isolated from infected spleens, and PCR was performed to evaluate the integrity of the *M2* locus. Cellular *GAPDH* serves as an amplification control. (D) Single-cell suspensions of spleen cells were serially diluted, and the frequencies of cells harboring MHV68 genomes were determined using a limiting-dilution PCR analysis. (E) Reactivation frequencies were determined by *ex vivo* plating of serially diluted cells on an indicator monolayer. Cytopathic effect was scored 2-3 weeks post-plating. Groups of 3–5 mice were pooled for each infection and analysis. Results are means of 2–3 independent infections. Error bars represent standard error of the means. ** denotes p<0.01 in a two-tailed student’s t-test.

### M2 is required in AID-expressing B cells for MHV68 latency and reactivation

During latency establishment, GHVs are thought to usurp GC reactions to gain access to the pool of long-lived MBCs. At early latency time points the majority of MHV68-infected cells exhibit a GC phenotype (2). M2 expression alone can drive naïve B cells toward a GC B cell phenotype and steer infected B cell differentiation toward a plasma cell phenotype (28, 29). Due to the direct role of M2 in B cell maturation processes and the importance of GC B cells during early latency time points, we hypothesized that M2 expression functions within GC B cells to promote latency establishment.

To determine how M2 functions within the GC B cell compartment, we evaluated M2.loxP infection in mice that express Cre recombinase under control of the promoter for the activation-induced cytidine deaminase (AID), the enzyme responsible for somatic hyper-mutation (SHM) and class-switch recombination (CSR) in GC B cells (19). Following IN infection of AID-Cre mice, M2.loxP viral titers in the lung were equivalent to WT MHV68 on day 7 post-infection (**Fig 4A**). A PCR analysis of the *M2* locus demonstrated that *M2* was deleted in these mice (**Fig 4C**). Similar to results in CD19-Cre mice, M2.loxP caused reduced splenomegaly relative to WT virus on day 16 post-infection (**Fig 4B** and **Table 1**), and M2.loxP latency in the spleen was 10-fold lower than WT MHV68 at this time point (**Fig 4D**). Additionally M2.loxP reactivation was below the limit of detection of 1 in 100,000 cells (**Fig 4E** and **Table 2**), which indicates that M2 expression in AID-expressing B cells potentiates MHV68 reactivation. These data strongly suggest that M2 expression within the GC compartment drives efficient latency establishment in and reactivation from the spleen following intranasal infection.

**Figure 4.**
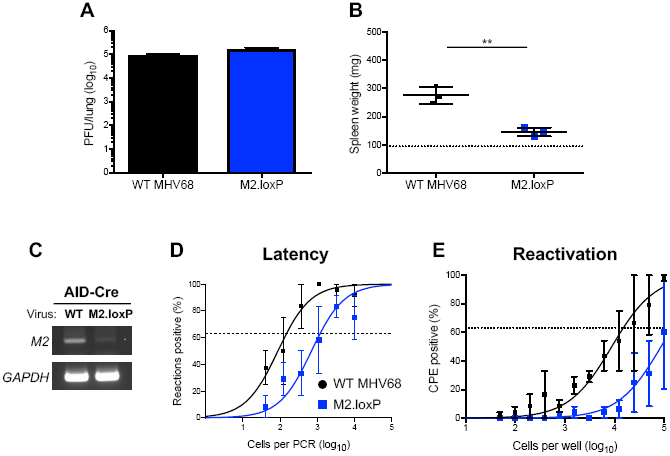
M2 deletion in AID-expressing cells impairs MHV68 latency and reactivation. AID-Cre mice were infected IN with 1000 PFU of the indicated viruses. (A) Mice were sacrificed on day 7 post-infection, and viral titers in lung homogenates were determined by plaque assay. (B-E) Mice were sacrificed on day 16 post-infection. (B) Spleens were harvested and weighed as a measure of splenomegaly. The dashed line indicates the average mass of spleens from mock-infected mice. Each dot represents one mouse. (C) DNA was isolated from infected spleens, and PCR was performed to evaluate the integrity of the *M2* locus. Cellular *GAPDH* serves as an amplification control. (D) Single-cell suspensions of spleen cells were serially diluted, and the frequencies of cells harboring MHV68 genomes were determined using a limiting-dilution PCR analysis. (E) Reactivation frequencies were determined by *ex vivo* plating of serially diluted cells on an indicator monolayer. Cytopathic effect was scored 2-3 weeks post-plating. Groups of 3–5 mice were pooled for each infection and analysis. Results are means of 2–3 independent infections. Error bars represent standard error of the means. ** denotes p<0.01 in a two-tailed student’s t-test.

### M2 is not required within GC B cells for colonization of lymphoid tissue

Following IN inoculation, MHV68 drains from the lungs to the mediastinal lymph nodes (MLNs) (32). From MLNs the virus disseminates via the blood to distal latency reservoirs, such as the spleen (33). We previously demonstrated that *ORF73*, the viral gene that encodes the MHV68 LANA homolog, is not necessary in CD19^+^ B cells for initial viral deposition in MLNs, despite its critical importance in hematogenous dissemination (30). Whether M2 is required for MLN infection and its requirement in specific B cell subsets has not been tested. We therefore evaluated MLN infection for M2.loxP, M2.Stop and WT MHV68 in either C57BL/6 or AID-Cre mice. In C57BL/6 mice, all three viruses were detected at equivalent levels in MLNs on day 16 after IN infection. In agreement with these findings, M2.loxP also established latency at levels similar to WT MHV68 in the MLNs of AID-Cre mice (**Fig 5A** and **5B**). These data support the conclusion that M2 is not necessary for latency in draining lymphoid tissue. Given that latency in the spleen is reduced in the absence of M2 following IN inoculation, we hypothesize that MHV68 facilitates viral dissemination to the spleen after initial seeding of the draining MLN.

**Figure 5.**
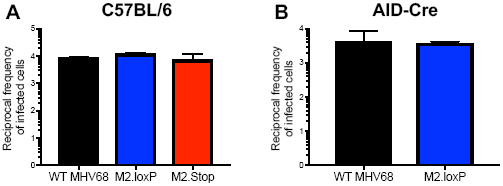
M2 expression is not required for MHV68 colonization of draining lymph nodes. (A) C57BL/6 or (B) AID-Cre mice were infected IN with 1000 PFU of indicated virus. Single-cell suspensions of MLNs were serially diluted and the frequencies of cells harboring MHV68 genomes were determined using a limiting-dilution PCR analysis. Reciprocal frequencies shown. Groups of 3–5 mice were pooled for each infection and analysis. Results are means of 2–3 independent infections. Error bars represent standard error of the means.

Intraperitoneal (IP) inoculation with MHV68 is thought to provide direct viral access to the spleen, bypassing any requirements for trafficking that may impact latency establishment following IN inoculation (27, 34). Similar to observations for MLNs, M2-null MHV68 achieves normal levels of latency in the spleen after IP inoculation, but still exhibits a reactivation defect (27, 28, 35). To determine how deletion of *M2* from AID-expressing B cells influenced chronic MHV68 infection after IP inoculation, we compared infections of M2.loxP and WT MHV68 in AID-Cre mice. Analogous to observations following IN inoculations, M2.loxP exhibited reduced splenomegaly compared to WT virus following IP infection of AID-Cre mice (**Fig 6A**). While the frequency of cells harboring latent M2.loxP genomes in spleens of AID-Cre mice was only slightly reduced compared to WT MHV68, M2.loxP was severely impaired in its capacity to reactivate from splenic latency (**Fig 6B** and **6C**). In peritoneal exudate cells (PECs), M2.loxP and WT MHV68 latency and reactivation were equivalent, which is consistent with the notion that MHV68 does not use AID-expressing cells to access or reactivate from cells present in the peritoneum (**Fig 6D** and **6E**). We also infected CD19-Cre mice as a control to confirm that phenotypes were similar when *M2* was deleted in B cells in general. Again, M2.loxP established latency in the spleen at levels similar to WT MHV68, but still exhibited a reduction in reactivation (**Fig 6F** and **6G**). Reduced detection of the *M2* locus in M2.loxP-infected AID-Cre and CD19-Cre mice indicates that *M2* was efficiently deleted from both strains after IP infection (**Fig 6H**). Together, these data suggest that M2 is not required for viral latency establishment within primary lymphatic tissues, but functions after deposition in draining lymph nodes to promote viral dissemination to the spleen.

**Figure 6.**
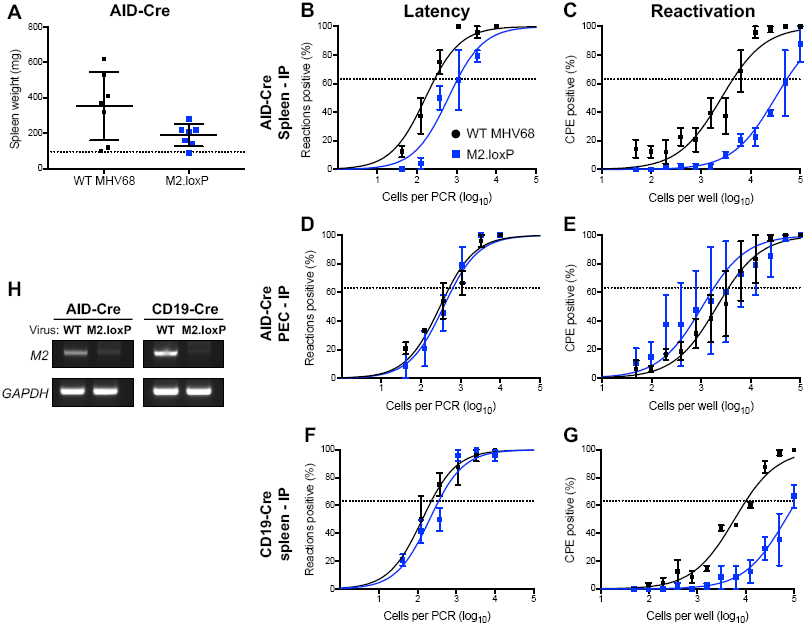
M2 expression in B cells is necessary for reactivation, but not latency establishment, following direct lymphoid tissue infection. AID-Cre (A-E, H) or CD19-Cre (F-H) mice were infected IP with 1000 PFU of the indicated virus and sacrificed on day 16 post-infection. (A) Spleens were harvested and weighed as a measure of splenomegaly. The dashed line indicates the average mass of spleens from mock-infected mice. Each dot represents one mouse. Single-cell suspensions of spleen cells (B, F) or PECs (D) were serially diluted, and the frequencies of cells harboring MHV68 genomes were determined using a limiting-dilution PCR analysis. Reactivation frequencies of splenocytes (C, G) or PECs (E) were determined by *ex vivo* plating of serially diluted cells on an indicator monolayer. Cytopathic effect was scored 2-3 weeks post-plating. (H) DNA was isolated from infected spleens of the indicated mouse strain, and PCR was performed to evaluate the integrity of the *M2* locus. Cellular *GAPDH* serves as an amplification control. Groups of 3–5 mice were pooled for each infection and analysis. Results are means of 2–3 independent infections. Error bars represent standard error of the means. ns denotes non-significant in a two-tailed student’s t-test. ** denotes p<0.01 in a two-tailed student’s t-test.

### M2 deletion from AID-expressing cells only moderately reduces plasmablast infection

M2-driven plasma cell differentiation accounts for ∼90% of MHV68 *ex vivo* reactivation (28). M2 expression in BCL-1 lymphoma cell lines promotes features of plasmacyte morphology, including increased size, granularity, and secretion of high levels of IgM. The notion that M2 drives plasmablast differentiation is consistent with reports that M2.Stop virus persists in B lymphocytes which have not undergone class-switching (IgD^+^IgG2a^-^) (28, 36). To determine whether M2 deletion from AID-expressing cells influences infection of plasmablasts in the spleen, we sorted CD138^+^/CD38^lo^ cells from spleens of infected AID-Cre mice (37) (**Fig 7A**) and performed qPCR to quantify viral genomes. Interestingly, by this method of quantification, we observed a 4.5 cycle difference, indicative of a ca. 20-fold deficit, in M2.loxP genome levels compared to WT virus in unsorted cells (**Fig 7B**). However, M2.loxP DNA in the plasmablast population was essentially equivalent to WT MHV68 (**Fig 7C**). We suspect that the larger impact on the general level of viral genomes detected in this experiment reflects that qPCR provides an absolute quantification of viral genomes in the population, rather than the evaluation of cells with at least one genome that is provided by the standard LD-PCR approach. This suggests that, while similar numbers of cells harbor MHV68 genomes (**Fig 6**), there are fewer viral genomes per cell upon deletion of *M2* from AID-expressing cells. In contrast, the absence of a severe phenotype for plasmablast infection suggests that infection of plasmablasts is either less dependent on M2 or does not require prior infection of an AID-expressing cell.

**Figure 7.**
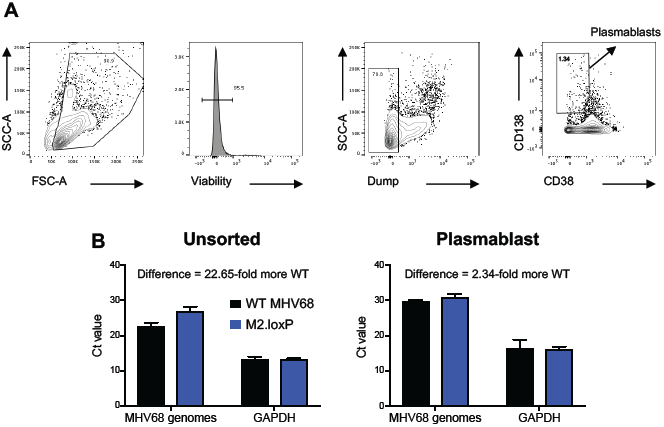
Loss of M2 in AID-expressing cells does not reduce infection of plasmablasts. AID-Cre mice were infected IP with 1000 PFU of the indicated virus and were sacrificed on day 14 post-infection. Spleens were harvested and stained as indicated for FACS-based sorting. Representative plots in panel (A) demonstrate the gating strategy used for sorting of CD138^+^CD38^lo^ plasmablasts. (B) DNA was isolated from unsorted splenocytes or sorted plasmablasts and qPCR was performed on 50 ng of DNA to detect MHV68 genomes or cellular *GAPDH* as a control. Fold difference in MHV68 genome quantities was determined using the ΔΔCt method.

## DISCUSSION

Gammaherpesviruses have evolved to potently manipulate and usurp immune responses generated toward viral antigens. For instance, studies in SAP- and IL-21 receptor-deficient mice demonstrate that MHV68 utilizes the general GC response to its own infection as a means to expand the latent cellular reservoir and generate long-lived reservoirs like MBCs (21, 38, 39). Understanding how specific viral genes engage these immune pathways is important for understanding the basic biology of GHV infection. Through the generation of null and point mutant viruses, overexpression studies, and utilization of fluorescent tagging, the MHV68 M2 protein is already linked to manipulation of B cell activation and differentiation (22, 27-29, 35); but when and where M2 exerts its functions to facilitate chronic viral infection and reactivation from latency are not known. To better define M2 functions in MHV68 pathogenesis, we describe here the generation and *in vivo* characterization of M2.loxP, a recombinant virus with the capacity to conditionally delete the *M2* locus in a Cre-dependent manner. Using this system, we demonstrate that loss of *M2* from AID-expressing B cells essentially recapitulates M2-null virus phenotypes in latency establishment and reactivation. We therefore conclude that M2 primarily functions within the GC B cell compartment to facilitate chronic MHV68 infection (**Fig 8**).

**Figure 8.**
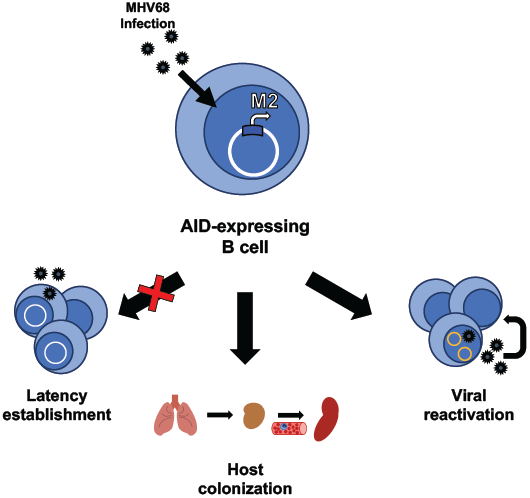
Model of cell-type-specific requirements for M2 in MHV68 latency. The majority of M2-null MHV68 phenotypes are recapitulated by specific *M2* deletion in AID-expressing B cells. These phenotypes overlap with those observed when *M2* is deleted in all CD19^+^ B cells. M2 is not directly required for viral infection of and apparent initial expansion in lymphoid tissues, but plays a role in facilitating dissemination within the host and reactivation during early latency time points. Results of plasma cell infection were inconclusive.

While our results indicate that the presence of *loxP* sites flanking *M2* in the MHV68 genome does not overtly influence lytic replication and latency, there are potential caveats to consider. The effect of *loxP* insertion on adjacent gene expression has not been assessed. The coding capacity of the MHV68 genome is densely packed (40, 41), and it is possible that *loxP* insertion or deletion of the locus inadvertently disrupts the function of overlapping or adjacent transcripts. To better characterize the potential off target effects that could occur following *M2* deletion, further analysis such as single-cell RNA sequencing is required. Additionally, if paired with mechanisms for marking infected cells, such as expression of a fluorescent or enzymatic reporter (2, 42), the impact of *M2* deletion on downstream infection of specific cell populations could be evaluated. Moreover, a marking approach could further enhance evaluations of M2 functions in signal transduction (24, 43-45) and impacts on cellular transcription (24), especially manipulation of the germinal center response. As mentioned earlier, M2-null MHV68 accumulates in naïve B cells during latency establishment and does not infect plasma cells (28, 36). Our qPCR data following sorting of plasmablast populations suggests that either MHV68 passage through AID-expressing cells only minimally influences plasmablast infection or that these cells are directly infected by the virus. Use of marking approach as described above to specifically isolate the infected cells could enhance attempts to further phenotype the infected plasmablast population, and infection of mice that express Cre in plasma cells could also be performed in future experiments.

MHV68 colonization of the host is thought to require viral dissemination from primary sites of infection into draining lymph nodes, initial expansion in the lymph node, and subsequent relay via the blood to secondary lymphoid organs, such as the spleen (11, 30, 32). Our data with both M2.Stop and M2.loxP demonstrate that M2 is not required for MHV68 to reach the draining lymph nodes. Similarly, using IP infection to bypass trafficking requirements for colonization of the spleen, both M2-null (27, 35, 36) and M2.loxP MHV68 efficiently establish latency in the spleen, though viral reactivation is severely impaired. These findings suggest that M2 is not directly required for MHV68 to establish latency in lymphoid tissue, but is required for downstream colonization events. Since M2.loxP accumulates to levels that are equivalent to WT virus in the MLN after IN infection of AID-Cre mice, but does not efficiently infect the spleen, it is reasonable to conclude that MHV68 traffics through a GC B cell to facilitate the seeding of distal latency reservoirs. Clearly this is not absolute, as viral genomes are still detected in the spleen following IN infection of Cre-expressing mice.

Since AID expression correlates with B cell maturation, we consider it unlikely that the M2.loxP defect corresponds to the accumulation of M2.Stop virus in naïve B cells, since M2 would presumably still be present in naïve B cells, due to a lack of AID-expression (46). M2-driven reactivation facilitating seeding of the spleen is possible, however another virus in which we conditionally delete *ORF50*, the gene that encodes the replication and transcription activator (RTA), suggests that reactivation/lytic replication in B cells is not required for splenic latency after IN inoculation (manuscript in preparation). We speculate that M2’s function in augmenting signaling within GC B cells is critical to enhancing MHV68 trafficking to secondary sites of latency in the mouse. We anticipate pairing cell-type-specific ablation of M2 function with single-cell transcriptomic and proteomic approaches in future experiments to evaluate these possibilities.

## MATERIALS AND METHODS

### Ethics statement

Mouse experiments were carried out in accordance with National Institutes of Health, United States Dept. of Agriculture, and UAMS Division of Laboratory Animal Medicine and Institutional Animal Care and Use Committee (IACUC) guidelines. The protocol supporting this study was approved by the University of Arkansas for Medical Science (UAMS) IACUC (animal use protocol 3817). Mice were anesthetized prior to inoculations and sacrificed humanely at the end of experiments.

### Cells and viruses

NIH 3T12 (ATCC CCL-164), 3T12 Flp+ (30), BHK21 (ATCC CCL-10), Vero (ATCC CCL-81), Vero-Cre (47), were cultured in Dulbecco’s Modified Eagle Medium (DMEM) supplemented with 10% fetal bovine serum (FBS), 100 U/ml penicillin, 100 ug/ml streptomycin, and 2 mM L-glutamine (cDMEM). Cells were maintained at 37°C in 5% CO_2_ and ∼99% humidity. Murine embryonic fibroblasts (MEFs) were harvested from C57BL/6 mouse embryos as previously described (34). Previously described viruses used in this study include FRT BAC MHV68 (WT MHV68) (30) and M2.null MHV68 (M2.STOP) (27).

To generate the M2.loxP BAC, *loxP* sites were inserted adjacent to the 5’ and 3’ ends of M2 in a FRT BAC template by two successive rounds of *en passant* mutagenesis as previously described (30) utilizing primers:

M2loxpUP_for

5’ – GTCTGTCACGCTTCTCCTTCCAGGCGTGTTTAAAGAAAAAATAACTTCGTATAGCATAC

ATTATACGAAGTTATGTTATGTTCTGCGTTAGCACTAGGGATAACAGGGTAATCGATTT – 3’

M2loxpUP_rev

5’ – GGTTAATTGGTTGTAACACTGGCCAGGCGTGTTTAAAGAAAAAATAACTTCGTATAGCATACATTATACGAAGTTATGTTATGTTCTGCGTTAGCACCTTCACTGTTACTCCTCGCC – 3’

M2loxpDWN_for

5’ – GTGGGGGTGTTGGGGCCATTACCTGAAAACGAAACCTCATATAACTTCGTATAGCATACATTATACGAAGTTATCATCAGACTAGTGCCTGTTGTAGGGATAACAGGGTAATCGATTT-3’

M2loxpDWN_rev

5’ – CCTTTTACTGGAGGGGGTTTCAACAGGCACTAGTCTGATGATAACTTCGTATAATGTATGCTATACGAAGTTATATGAGGTTTCGTTTTCAGGTGCCAGTGTTACAACCAATTAACC – 3’.

Viruses were passaged in Flp-expressing 3T12 fibroblasts to remove the BAC cassette, and titers were quantified as previously described (30).

### Mouse infections and tissue harvests

Male and female, C57BL/6, CD19-Cre (B6.129P2(C)-Cd19tm1(cre)Cgn/J), and AID-Cre (B6.129P2-Aicdatm1(cre)Mnz/J) were purchased from Jackson Laboratories. Mice were bred and maintained according to all local, state, and federal guidelines under the supervision of the UAMS Division of Laboratory Animal Medicine. Eight-to-ten-week-old mice were anesthetized using isoflurane and inoculated with 1000 PFU of viruses diluted in incomplete DMEM (20 μl) for IN inoculations or injected with 1000 PFU of virus diluted in incomplete DMEM (200 ul) for IP inoculations. Splenocytes and peritoneal exudate cells (PECs) were harvested as previously described (48). Cells from draining lymph nodes were isolated as previously described (30).

### Splenocyte isolation and limiting-dilution analyses

Spleens were homogenized in a tenBroek tissue disrupter. Red blood cells were lysed by incubating tissue homogenate in 8.3 g/L ammonium chloride for 10 minutes at room temperature with shaking. Cells were filtered through 40 μM mesh to reduce clumping. Frequencies of cells harboring MHV68 genomes were determined using LD-PCR analysis as previously described (49). Frequencies of latently-infected cells capable of reactivating were determined using a limiting-dilution analysis for cytopathic effect induced on an indicator BHK21 monolayer as previously described (34).

### Plaque Assays

Plaque assays were performed as previously described using BHK21 cells (2 × 10^5^ cells/well). Briefly, infected cells were overlaid with 1.5% methylcellulose in DMEM supplemented with 2.5% calf serum, 100 U/ml penicillin, 100 μg/ml streptomycin, and 2 mM L-glutamine, and incubated at 37°C for 4-6 days. Cell monolayers were stained with a solution of crystal violet in formalin for identification and quantification of plaques.

### Cell Sorting

Splenocytes were isolated and sorted by flow cytometry. Blocking and detection antibodies were diluted in PBS with BSA and EDTA. Splenocytes were blocked with anti-CD16/32 (BD Biosciences) for 15 mins at 4°C prior to surface staining for 30 mins in the dark at 4°C. Dead cells were labeled using Fixable Viability Dye eFlour 780 (eBioscience) as per manufacturer’s instructions. Surface stains include: dump gate (CD3, Ter119, CD11b, CD11c-PerCp5.5), B220-redFluor™ 710 (Tonbo), CD19-BV650, CD38-PE-Cy7, CD138-PE (Biolegend). Cells were sorted on a FACS Aria using FACS Diva software. Data was analyzed using FlowJo software (v10.6.2).

### Nucleic acid isolation and qPCR

DNA was isolated by phenol/chloroform extraction and ethanol precipitation. 50 ng of DNA was diluted in PowerUp SYBR Green Master Mix (ThermoFisher) and analyzed by quantitative real-time PCR in an Applied Biosystems QuantStudio thermocycler. Cycling conditions were 10 min at 95°C followed by 40 cycles of 15 s at 95°C and 1 min at 60°C, using primers specific to the viral *ORF72* gene or cellular *GAPDH* gene (3, 50). Biological triplicate samples were analyzed in technical duplicate using the ΔΔ*CT* comparative threshold cycle method as previously described (50). Fold change in *ORF72* levels for M2.loxP was calculated relative to WT MHV68.

## ACKNOWLEDGEMENTS

We thank the laboratory of Dr. Jason Stumhofer for assistance in flow cytometry antibody panel design and Andrea Harris in the UAMS Flow Cytometry Core. This work was supported in part by R01CA167065 of the NIH National Cancer Institute and start-up funds from the UAMS College of Medicine and Arkansas Biosciences Institute to J.C.F. The Flow Cytometry Core and work described here also was supported in part by the Center for Microbial Pathogenesis and Host Inflammatory Responses award P20GM103625 from the NIH National Institute of General Medical Sciences Centers of Biomedical Research Excellence. D.W.W. and D.G.O. were supported by the Gundersen Medical Foundation. The funders had no role in study design, data collection and interpretation, or the decision to submit the work for publication.

